# Vaccination can drive an increase in frequencies of antibiotic resistance among non-vaccine serotypes of *Streptococcus pneumoniae*

**DOI:** 10.1101/135863

**Authors:** Uri Obolski, José Lourenço, Sunetra Gupta

## Abstract

The bacterial pathogen Streptococcus pneumoniae is a major public health concern, being responsible for more than 1.5 million deaths annually through pneumonia, meningitis and septicemia. In spite of vaccination efforts, pneumococcal carriage and disease remain high, since available vaccines target only a subset of serotypes and vaccination is often accompanied by a rise in non-vaccine serotypes. Epidemiological studies suggest that such a change in serotype frequencies is often coupled with an increase of antibiotic resistance among non-vaccine serotypes. Building on previous multi-locus models for bacterial pathogen population structure, we have developed a theoretical framework incorporating variation in serotype and antibiotic resistance to examine how their associations may be affected by vaccination. Using this framework, we find that vaccination can result in rapid increase in frequency of pre-existing resistant variants of non-vaccine serotypes due to the removal of competition from vaccine serotypes.

## Introduction

The bacterial pathogen, Streptococcus pneumoniae (or the pneumococcus) is estimated to be responsible for a third of all pneumonia cases and annually causing tens of millions of severe infections worldwide [1]. Two major tools are available for reducing pneumococcal disease: antibiotic treatment, and vaccination [2]. Antibiotic drugs remain an efficient way of clearing pneumococcal infections, but their efficiency is often impaired by the emergence and spread of antibiotic resistant pneumococci [3]. Vaccination can prevent new pneumococcal infections by increasing host immunity against pneumococcal serotypes, and eventually substantially reduce the incidence of infection by creating herd immunity [4, 5]. However, available vaccines, such as the pneumococcal conjugate vaccine (PCV), target only a subset of circulating pneumococcal serotypes, hence exerting selective pressure which can shift serotype frequencies – a phenomenon termed Vaccine induced Serotype Replacement (VISR) [6-8]. Changes in genetic composition of pneumococci have also been observed after vaccination [9, 10], and it has been proposed that vaccination may induce a shift in metabolic profiles of non-vaccine strains (known as Vaccine Induced Metabolic Shift, or VIMS), as a consequence of resource competition amongst bacteria sharing the same metabolic alleles [11, 12].

The deployment of pneumococcal vaccines has also led to significant changes in antibiotic resistance frequencies. It is unsurprising that vaccines targeting pneumococcal serotypes that have high resistance frequencies would lead to the reduction of resistance at a population level [13-17]. However it is not easy to account for a post-vaccination increase in resistance frequencies of subsets of non-vaccine type (nVT) pneumococci, as has been repeatedly observed in a range of locations [18-26]. Interestingly, resistance frequencies have not changed equally between all serotypes; and, within serotypes, resistance to different antibiotics has not changed uniformly [18-26]. Figure 1A shows changes in antibiotic resistance within the 19A serotype, an nVT of PCV7, following the introduction of this vaccine in 2000 within pneumococcal isolates of children ≤ 5 years collected in Massachusetts [10]; we observe an increase in the minimum inhibitory concentration (MIC) to erythromycin between 2001 and 2007, whereas resistance to penicillin and ceftriaxone remains unchanged (Figure 1A). It should be noted that the increase in erythromycin resistance occurred in spite of a decline of antibiotic prescription in the population over the study period [27].

**Figure 1:**
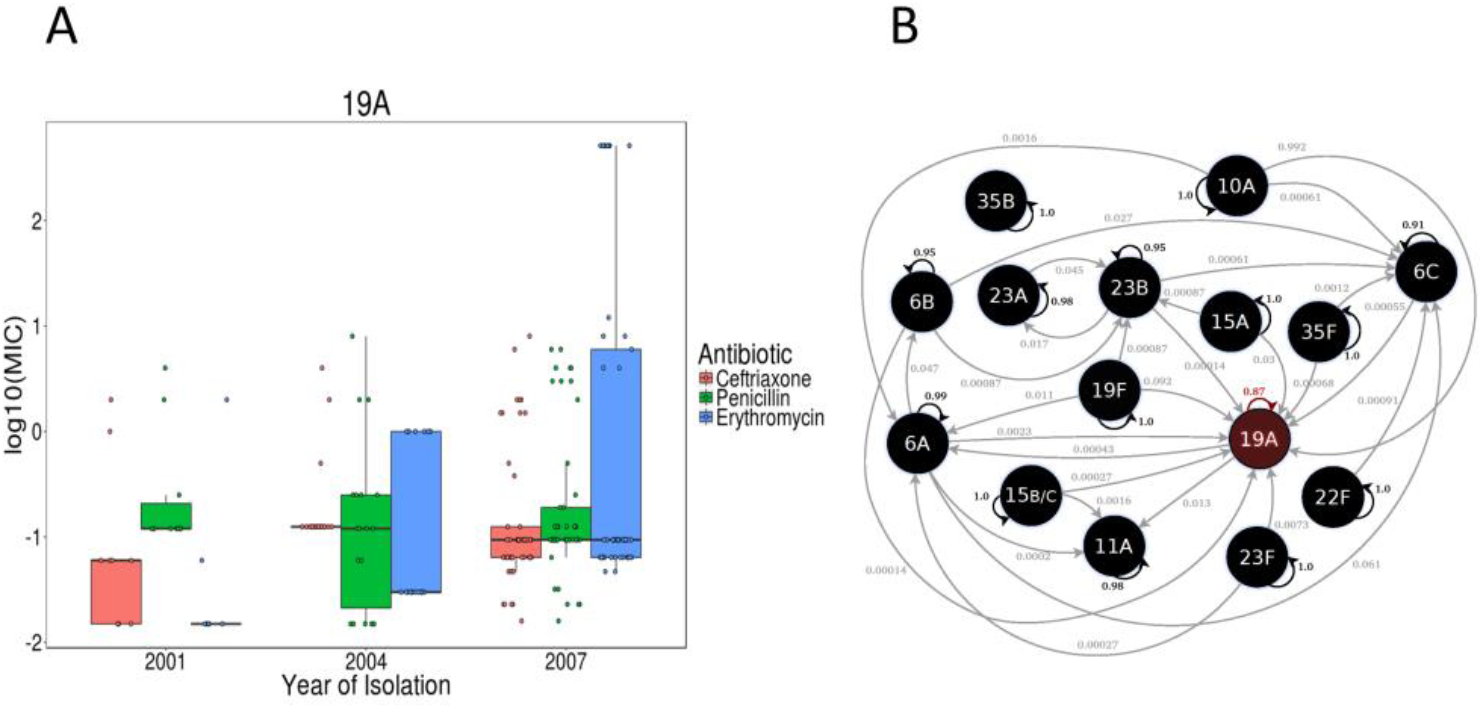
Observed changes in resistance, and admixture estimation of the 19A serotype. **(A)** 19A increases in MIC to erythromycin (p-value< 2 × 10^−7^, Spearman correlation= 0.58), but not in penicillin and ceftriaxone unchanged (p-value= 0.56, 0.57; Spearman correlation= -0.07;both), between 2001 and 2007 after PCV7 introduction at 2000. **(B)** Admixture analysis using BAPS (see Methods) on antibiotic resistance-associated genes on the same data as in (A). Arrows represent statistically significant admixture events (p-value for each event <0.001), with arrow directions defining origin and destination of admixed alleles, and numbers representing the fraction of alleles contributed from source to receiving serotype. 19A has the highest fraction of alleles most likely from other serotypes, with only 87% of alleles likely to originate from the serotype’s population.

A Bayesian Analysis of Population Structure (BAPS) revealed significant admixture of resistance-associated alleles from various different serotypes (Figure 1B; see Methods for full description). This analysis showed that only 87% of antibiotic resistance-associated allele combinations within serotype 19A were found to have originated from the 19A serotype itself, with the rest much more likely to be found in other serotypes. This exemplifies that 19A strains, notorious for increasing in resistance post-vaccination [18, 28-30], may experience changing selective pressures leading to changes in their distribution post-vaccination. The analysis was repeated for another publicly available UK pneumococcal data set [31] wherein the 19A serotype had the second highest value of antibiotic resistance alleles most likely originating from other serotypes (Supplementary Information Figure S1).

Here we explore the interactive evolution of vaccination and antibiotic resistance using a multi-locus model of serotype and antibiotic resistance with full flexibility in associations between alleles, reflecting the high levels of admixture shown above. In contrast with previous multi-locus models of pneumococcal evolution [11, 12] which assume interference occurs between organisms carrying similar metabolic and virulence alleles, we assume that antibiotic resistant strains are less likely to co-infect individuals infected with a susceptible strain due to ecological competition. We find that a post-vaccination surge in antibiotic resistance frequencies can occur in nVTs under these circumstances and, furthermore, may be hastened by asymmetries in rates of acquisition of resistance to different antibiotics.

## Results

### Model structure

We investigate the impact of vaccination within a system containing two streptococcal serotypes, *a* and *b*, of which the first is included in the vaccine (VT) and the second is not (nVT). We assume that immunity is serotype-specific, but may be incomplete, with its efficacy represented in our model by the parameter 0≤ γ ≤1, where γ=1 implies that immunity is complete and γ=0 corresponds to no serotype-specific immunity. We denote a bacterial strain of serotype *a* or *b* as having a resistance profile *j*, where *j* takes values in {00,01,10}, corresponding to sensitivity to both antibiotics, resistance to antibiotic *X*, and resistance to antibiotic *Y*.

The intrinsic transmissibility of a strain can be represented by its basic reproductive number [32], R_0_, which is a product of the duration of infection (D) and degree of infectivity (β), where the latter is effectively a combination of parameters defining the likelihood of acquisition by a susceptible individual of a particular strain from an infected individual. We assume that the cost of resistance would typically translate into lower infectivity of resistant strains compared to sensitive strains (β_00_ > β _01_); however, the duration of carriage may be longer for resistant strains due to antibiotic usage (D_01_>D_00_). Thus, in the absence of antibiotic usage, the basic reproductive number of resistant strains, R^01^_0_, will typically be lower than the than the basic reproductive number of sensitive strains, R^00^_0_, but this can be reversed with antibiotic usage. We further assume that intrinsic fitness differences (such as in growth rates) between resistant and susceptible strains may allow an individual carrying a susceptible strain of pneumococci to suppress co-infection by a resistant strain to a degree 0≤ ψ ≤1. Note that this is a form of ecological competition between bacterial strains and is not mediated by immunity: thus individuals carrying strain *a*_*00*_ may not be available for co-infection by either *a*_*01*_ or *b*_*01*_(for example if ψ = 1) but will be fully susceptible to further infection by *b*_*00*_.

### Vaccination can increase the frequency of antibiotic resistance among nVTs

We start by considering a model in which the two serotypes, *a* and *b*, are either susceptible or resistant to a single antibiotic (Y) and there are no serotype-specific differences in R_0_. The equilibrium frequencies of the four strains (*a*_*00,*_ *a*_*01*_, *b*_*00*_*, b*_*01*_) before vaccination are determined by the degree of serotype-specific immunity (γ) and inhibition of co-infection by resistant strains (ψ) within the system, and the basic reproductive numbers of sensitive and resistant strains (R^s^_0_ and R^r^ respectively). High serotype-specific immunity leads to the competitive exclusion within a given serotype, typically of the strain with the lower R_0_ [33]; however, the inhibition of co-infection by susceptible strains (ψ > 0) places a further cost on resistant strains such that they may be excluded even if they have a slightly higher R_0_. Under these circumstances, the removal of serotype *a* (the VT) through vaccination may cause a reversal of the outcome of competition between *b*_*00*_ and *b*_*01*_, with the resistant nVT completely replacing the susceptible nVT, as shown in Fig 2A. This is due to the removal of ecological competition between *a*_*00*_ and *b*_*01*_, and can be observed under circumstances where R^01^ is somewhat in excess of R^00^, provided ψ is above the following threshold (see Supplementary Information S2 for derivation):

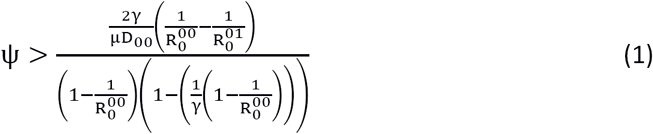

**Figure 2:**
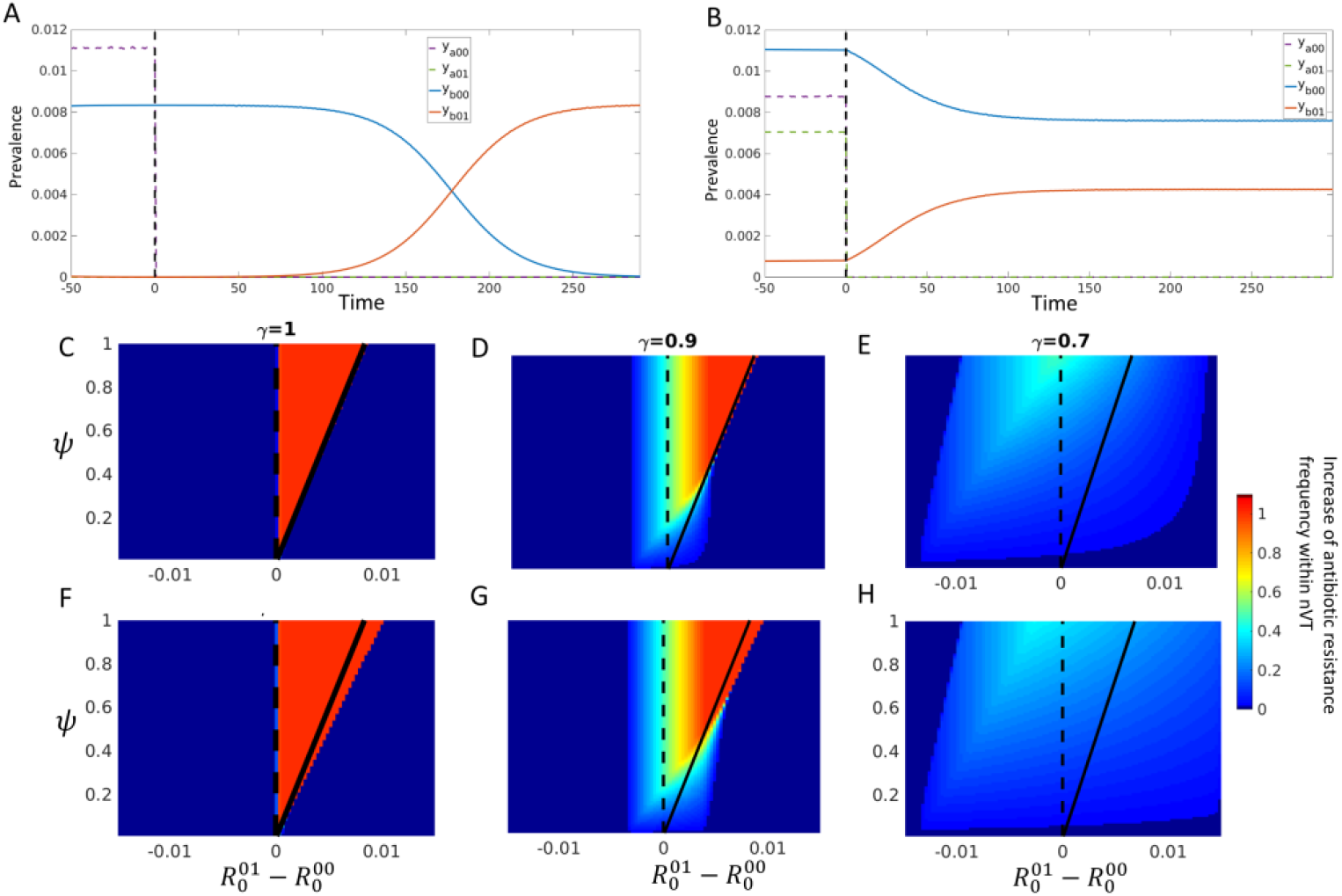
Vaccination can increase frequency of antibiotic resistance among non-vaccine serotypes. **(A)** Dynamics of susceptible VT (dashed purple), resistant VT (dashed green), susceptible nVT (blue) and resistant nVT (orange) pre- and post-vaccination (the time of which is marked by the dashed vertical line) under full serotype-specific immunity (γ=1, ψ =0.8; R^00^_0_=2.008; R^00^_0_=2) **(B)** Dynamics of the same model under intermediate serotype-specific immunity (γ=0.7,*ψ* =0.8; R^00^_0_=1.998; R^00^_0_=2). **(C-E)** Heat maps of the increase in relative frequency of the resistant nVT (*b01*) post-vaccination, against varying values of co-infection inhibition (Ψ), and the difference in reproductive number between resistant and susceptible nVTs (R^r^_0_-R^s^_0_, with R^s^_0_=2).The black curves are derived from eqn (1) in the main text, and mark the parameter ranges under which *b01* switches from being the rare to the common type post vaccination. **(F-H)** VTs are assigned increased transmission, expanding the range of parameters whereas surge in resistance frequencies occurs; the curves obtained from eqn(1) have been overlaid for ease of comparison with C-E. For A,B and F-H the transmission of the VTs was increased to1.5 fold of the nVTs reproductive number. Other parameters: D=1/ *σ* =30 days, *μ* =1/5 years^−1^.

When serotype-specific immunity is complete (γ = 1), the region where a surge in frequency of the resistant nVT occurs is limited to the area between R^r^_0_ > R^s^_0_ and the curve described by eqn (1). When R^01^_0_ <R^00^_0_, there is no change in the outcome of within-serotype competition as the susceptible NVT will continue to dominate after the VT is removed (Figure 2C, left of the dashed line); when R^01^_0_ is sufficiently in excess of R^00^_0_, *b*_*01*_ will have displaced *b*_*00*_ prior to vaccination and no change will be seen (Figure 2C, right of the solid black curve).

At lower levels of serotype-specific immunity (γ < 1), susceptible and resistant strains may coexist within the same serotype within certain boundaries of difference in R_0_: under these circumstances, vaccination can cause a surge in the frequency of the resistant nVT even when R^01^_0_ <R^00^_0_ (Figure 2 D&E) since this does not invariably lead to the total exclusion of *b*_*01*_. In this case, the resistant strain remains the rarer strain post-vaccination but may substantially increase in frequency, as shown in Figure 2B. In the region R^01^_0_>R^00^_0_, eqn (1) still determines whether the resistant nVT will increase from being the rarer strain before vaccination to being the more common strain following vaccination (Supplementary Information 2). To the right of the curve, *b*_*01*_ is already the more frequent strain before vaccination but may increase in prevalence after vaccination due to the cessation of competition from *a*_*00*_ (Figure 2 D&E).

Increasing the R_0_ of serotype *a* leads to an expansion in the parameter range within which there is a post-vaccination surge in *b*_*01*_ (Figure. 2 F-H). This is because the prevalence of *a*_*00*_ increases, causing *b*_*01*_ to be suppressed further; thus *b*_*01*_ experiences a greater increase in frequency when *a*_*00*_ is removed by vaccination Importantly, the general principles illustrated above remain unaltered when we introduce a second antibiotic, X, with pre-vaccination equilibria falling into 3 categories (i) coexistence of all strains (ii) competitive exclusion of both antibiotic resistant strains (b_10_ and b_01_) (iii) competitive exclusion of the resistant strain with the lower R_0_ (say b_10_). Following the removal of serotype *a* through vaccination, resistant strains that are already present will increase in frequency and there may be an emergence of strains that were excluded in the pre-vaccine era (Figure S2).

### Effect of asymmetries in rates of acquisition of resistance to different antibiotics

The model can be extended to explicitly incorporate rates of acquisition of resistance to two antibiotics, X and Y, by introducing the parameter ω_j_ to describe the probability of a sensitive strain acquiring a resistance profile *j*. We find that this has a significant impact on the outcome of vaccination where the less transmissible strain (in this case, b_10_, which is resistant to X) is associated with a higher rate of resistance acquisition (ω_01_ <ω_10_,). Under these circumstances, *b*_*10*_ may stabilise at higher pre-vaccination frequencies to *b*_*01*_, despite a significant transmissibility disadvantage (in Fig 3A, R_0_^10^=0.91R_0_^01^). The removal of competition from *a*_*00*_ due to vaccination effectively unmasks the transmission advantage of *b*_*01*_, thereby driving a rapid increase in its frequency. A stochastic implementation of this model indicates that a very significant rise in frequency of *b*_*01*_ can occur within a decade or two after the removal of the VT (Figure 3B) under realistic parameter combinations, and that it has a strong likelihood of eventually displacing *b*_*00*_ as the dominant strain within this serotype. Moreover, the time point when *b01* first becomes more common than *b10* is likely to be even within a decade from vaccination, but it is possible for it to occur after more than 50 years (Figure 1C). Therefore, substantial variance in surges of antibiotic resistant nVT strains post-vaccination is expected even between populations experiencing similar conditions.

**Figure 3:**
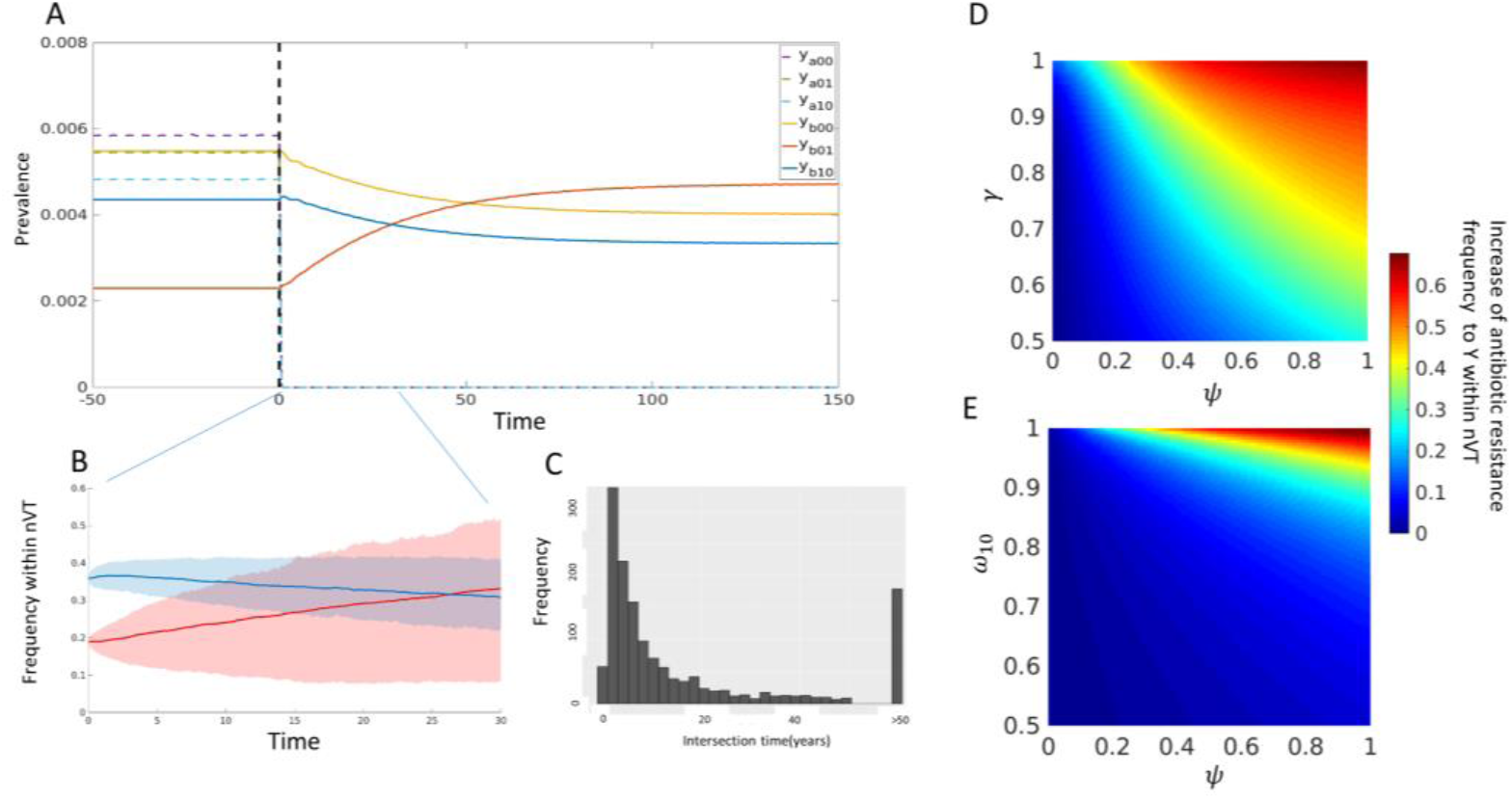
Increase in antibiotic resistance under asymmetric acquisition rates. **(A)** Dynamics of VT (solid curves) and nVT (dashed curves) pre- and post-vaccination, the time of which is marked by the dashed vertical line. We assume acquisition of resistance to X (b_00_ →b_10_) is higher than to Y (b_00_ →b_01_), but that b_10_ is less transmissible than b_01_. **(B)** Stochastic implementation of the post-vaccination dynamics (corresponding dynamics in (A) marked by the blue rectangle). Average values of 1500 simulations are given by solid lines, interquartile range of simulation results is given by shaded areas. **(C)** Histogram of the first time *b01* increases in frequency above *b10*, recorded from the 1500 simulations. Approximately 12% of simulations did not have an intersection between y_b01_ and y_b10_ during the simulated 50 years (marked in the histogram by the >50 bar). **(D,E)** Heat maps of the increase in relative frequency of b_01_ following vaccination (R_0_^10^=1.88, R_0_^00^= R_0_^01^=2; R_0_(a^j^)= 1.5 R_0_(b^j^); γ=0.7, ψ=0.9 and *ω*^10^ = 1 - *ω*^01^ = 0.99 when they are not varied).

As might be expected, increasing ω_10_ lowers the pre-vaccine frequency of *b*_*01*_, leading to a higher post-vaccination surge (Figure 3D). The increase in *b*_*01*_ is also more pronounced at higher values of ecological interference from *a*_*00*_ (ψ) (Figure 3 D & E), in line with our previous results. The increase in resistance among nVTs also depends on the strength of serotype-specific immunity (γ), as this determines the extent to which *b*_*01*_ can realise its transmission advantage (Figure 3E).

## Discussion

Understanding the population dynamics of *Streptococcus Pneumoniae* is an important endeavour from a public health perspective, and the post-vaccination surge in antibiotic resistance frequencies observed in some nVTS is of special concern.

Existing models of antibiotic resistance typically aim to define the conditions minimizing resistance emergence or spread under different antibiotic regimes [34-37]. Fewer efforts have been made to study the effects of pneumococcal vaccination on the evolution of antibiotic resistance [38-40] and only in one of these (Lehtinen et al [40]), as far as we are aware, has a mechanism been proposed by which vaccination may induce an increase in antibiotic resistance.

A crucial difference between our model and that of Lehtinen et al, is that we include serotype-specific immunity following natural infection, although by no means does this have to be completely sterilising. Indeed, stable coexistence of resistant and susceptible strains in the pre-vaccine era is more likely to occur, in our model, under incomplete immunity. In the absence of serotype-specific immunity, coexistence becomes difficult to obtain: Lehtinen et al. show, however, that heterogeneity in duration of carriage can maintain coexistence. Within their framework, the removal of vaccine strains permits longer duration of carriage, thereby leading to an increase in antibiotic resistance provided the associated genes are in epistasis with genes influencing carriage duration; Lehtinen et al. provide evidence implying such an epistatic interaction by analysis of carriage duration and antibiotic resistance from observed data.

Our model also relies on the removal of ecological interference from vaccine strains, but here this alters the outcome of competition between resistant and sensitive strains within an nVT, potentially leading to a surge in frequency of the resistant nVT. This provides a possible explanation to the surge in antibiotic resistance in nVTs, observed independently in various populations [18-26]. We account for scenarios facilitating a surge in frequency of all or only a subset of the resistant types, under coexistence or competitive exclusion, and for varying transmissibility of the different strains. Furthermore, we introduce the notion that a resistant nVT can increase even more in frequency following vaccination if it were masked by a low rate of acquisition of resistance to other antibiotics before vaccination.

We believe that further genetic and phenotypic data of pneumococci, pre- and post-vaccination, would help distinguish between these hypotheses, and eventually lead to the design of interventions that will prevent post-vaccination increases in antibiotic resistance.

## Methods

### Model

In our model, each strain genotype is defined by the tuple (*i*, *j*), where *i* determines serotype and *j* the antibiotic resistance allele, respectively. For the simple bi-allelic, two-locus case, let *i* ∈ (*a*, *b*), *j* ∈ (00,01). We denote by *y*_*ij*_ the proportion of individuals currently infected by strain *ij*; *z*_*i*_ is the proportion of the population previously exposed to serotype *i*; *Z*_*i*_ is the proportion of the population previously **or** currently exposed to serotype i; *Y*_*ij*_ and *V*_*ij*_ will refer to primary and secondary infections with strain *ij*, respectively. For example, the proportion of individuals infected by susceptible bacteria of serotype *a* is *y*_*a*00_; the proportion individuals previously exposed to serotype *a* is given by *z*_*a*_.

Let *z*_*a*_ be the proportion of individuals who have been infected with antigenic type *a*, and *y*_*a*01_ contain all individuals currently infected with *a*_01_. Let us first assume (i) that hosts infected by a bacterial strain *i*, *j* cannot be re-infected by a strain with the same serotype, *i* (ii) a host infected by bacteria susceptible to antibiotics can only be co-infected by susceptible strains of bacteria.

The equations of the epidemiological model are given by (full derivations are presented in Supplementary file S1):

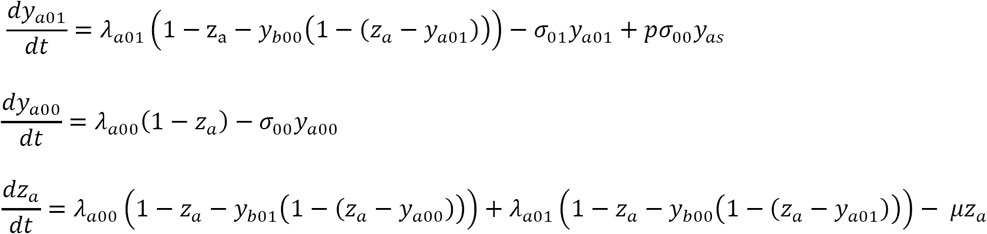

Where *σ*_*ij*_ is the rate of clearance, equivalent to the inverse of infection length *D*_*ij*_; *μ* is the host removal rate; *λ*_*ij*_ is the force of infection, determined by *y*_*ij*_ *β*_*ij*_, where *β*_*ij*_is the transmission rate of strain *ij*. An analogous set of equations is given for bacteria of serotype *b.* The basic reproduction number for strain *ij* is determined by 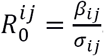
. Antibiotic resistance acquisition is also possible in our framework and will be denoted by the parameter *p*, determining the fraction patients acquiring antibiotic resistance instead of being cleared. When more than one locus determining antibiotic resistance is modelled, we will denote by ω^*l*^ the probability of acquiring resistance to a certain profile *l* by the susceptible strain. When not varied, parameter values are set to 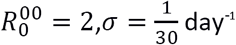
and 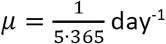
, in accordance with paediatric pneumococcal colonization [38]; *p* = 0.05 for resistance acquisition scenarios [41].

To relax the two assumptions introduced above, we introduce two parameters:

We represent serotype-specific immunity by 0 ≤ *γ* ≤ 1, where *γ* = 1 is equivalent to the assumption postulated above with complete specific immunity, and *γ* = 0 corresponds to no serotype-specific immunity.

Similarly, we introduce the parameter 0 ≤ Ψ ≤ 1 to represent the probability that an individual carrying a susceptible strain of pneumococci will suppress co-infection by a resistant strain, due to the fitness cost of antibiotic resistance.

Adding these parameters yields the new set of equations:

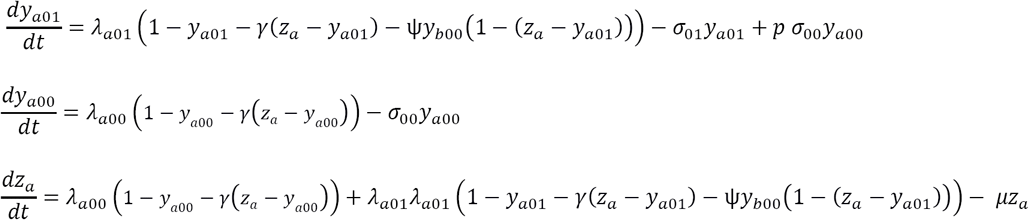

We can extend this model to any number of antigenic alleles, and any number of bi-allelic resistance loci (see Supplementary file S1). Vaccination is added to the model by reducing the *R*_0_ of vaccine strains by 90%.

### Stochastic implementation

We developed a semi individual-based implementation of our equations, based on the Gillespie stochastic simulation algorithm (SSA) [42]. Variables representing infected hosts (*y*_*ij*_) are explicitly modelled under the SSA framework, whereas previously infected individuals are approximated via a deterministic approach: at each newly drawn time point *t*_*n*+1_, the individuals previously infected with strain *i* are given by 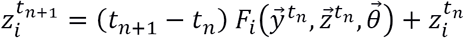

Where *F*_*i*_ marks the differential equation defined for the deterministic dynamics of patients previously infected with 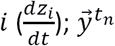
and 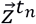
represent all values of currently and previously infected individuals at time *t*_*n*_; and 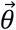
represents all parameters defined in the deterministic model. Since the number of previously infected patients is much greater than those currently infected, it can be approximated with a deterministic equation (updated by random processes). All simulations performed with a population size of 100,000. Finally, we constrain the number of hosts infected by any of the different strains to be ≥ 1, to avoid the absorbing states of strain extinction.

### Bayesian analysis of population structure (BAPS)

Two data sets of pneumococcal genomes, collected from the USA [10] and the UK [31], were annotated using the BIGSdb software and assigned alleles with the Genome Comparator tool (with ATCC 700669 pneumococcal strain as the reference genome) [43]. We examined 34 loci associated with antibiotic resistance (given in Table S1) and preformed an admixture analysis on predefined clusters based on serotypes in the BAPS 6 software [44, 45]. Only serotypes containing 15 or more samples were used, leaving us with N=514 and N=391 observations for the USA and UK data respectively. Parameters used in the software were set according to the higher accuracy recommendations given in the BAPS manual: max clusters – 50; iterations – 500; reference individuals – 200; iterations per reference individuals – 20. Admixture inclusion threshold was set to p-value <0.001.

## References

1. Organization, W.H., Measuring impact of Streptococcus pneumoniae and Haemophilus influenzae type b conjugate vaccination. 2012.

2. Parks, T., L. Barrett, and N. Jones, Invasive streptococcal disease: a review for clinicians. British medical bulletin, 2015. 115(1): p. 77–89.

3. Ventola, C.L., The antibiotic resistance crisis: part 1: causes and threats. Pharmacy and Therapeutics, 2015. 40(4): p. 277.

4. Kristiansen, P.A., et al., Impact of the serogroup A meningococcal conjugate vaccine, MenAfriVac, on carriage and herd immunity. Clinical infectious diseases, 2012: p. cis892.

5. Maiden, M.C., et al., Impact of meningococcal serogroup C conjugate vaccines on carriage and herd immunity. Journal of Infectious Diseases, 2008. 197(5): p. 737–743.

6. Hanage, W., Serotype replacement in invasive pneumococcal disease: where do we go from here? Journal of Infectious Diseases, 2007. 196(9): p. 1282–1284.

7. Weinberger, D.M., R. Malley, and M. Lipsitch, Serotype replacement in disease after pneumococcal vaccination. The Lancet, 2011. 378(9807): p. 1962–1973.

8. Lipsitch, M., Bacterial vaccines and serotype replacement: lessons from Haemophilus influenzae and prospects for Streptococcus pneumoniae. Emerging infectious diseases, 1999. 5(3): p. 336.

9. Beall, B.W., et al., Shifting genetic structure of invasive serotype 19A pneumococci in the United States. Journal of Infectious Diseases, 2011: p. jir052.

10. Croucher, N.J., et al., Population genomics of post-vaccine changes in pneumococcal epidemiology. Nature genetics, 2013. 45(6): p. 656–663.

11. Gupta, S., N.M. Ferguson, and R.M. Anderson, Vaccination and the population structure of antigenically diverse pathogens that exchange genetic material. Proceedings of the Royal Society of London B: Biological Sciences, 1997. 264(1387): p. 1435–1443.

12. Watkins, E.R., et al., Vaccination Drives Changes in Metabolic and Virulence Profiles of Streptococcus pneumoniae. PLoS Pathogens, 2015. 11(7).

13. Dagan, R., et al., Reduction of pneumococcal nasopharyngeal carriage in early infancy after immunization with tetravalent pneumococcal vaccines conjugated to either tetanus toxoid or diphtheria toxoid. The Pediatric infectious disease journal, 1997. 16(11): p. 1060–1064.

14. Dagan, R., et al., Effect of a nonavalent conjugate vaccine on carriage of antibiotic-resistant Streptococcus pneumoniae in day-care centers. The Pediatric infectious disease journal, 2003. 22(6): p. 532–539.

15. Mbelle, N., et al., Immunogenicity and impact on nasopharyngeal carriage of a nonavalent pneumococcal conjugate vaccine. Journal of Infectious Diseases, 1999. 180(4): p. 1171–1176.

16. Mishra, R.P., et al., Vaccines and antibiotic resistance. Current opinion in microbiology, 2012. 15(5): p. 596–602.

17. Chiba, N., et al., Rapid decrease of 7-valent conjugate vaccine coverage for invasive pneumococcal diseases in pediatric patients in Japan. Microbial Drug Resistance, 2013. 19(4): p. 308–315.

18. Moore, M.R., et al., Population snapshot of emergent Streptococcus pneumoniae serotype 19A in the United States, 2005. Journal of Infectious Diseases, 2008. 197(7): p. 1016–1027.

19. Hanage, W.P., et al., Diversity and antibiotic resistance among nonvaccine serotypes of Streptococcus pneumoniae carriage isolates in the post-heptavalent conjugate vaccine era. Journal of Infectious Diseases, 2007. 195(3): p. 347–352.

20. Kyaw, M.H., et al., Effect of introduction of the pneumococcal conjugate vaccine on drug-resistant Streptococcus pneumoniae. New England Journal of Medicine, 2006. 354(14): p. 1455–1463.

21. Park, S.Y., et al., Impact of conjugate vaccine on transmission of antimicrobial-resistant Streptococcus pneumoniae among Alaskan children. The Pediatric infectious disease journal, 2008. 27(4): p. 335–340.

22. Huang, S.S., et al., Post-PCV7 changes in colonizing pneumococcal serotypes in 16 Massachusetts communities, 2001 and 2004. Pediatrics, 2005. 116(3): p. e408–e413.

23. Pillai, D.R., et al., Genome-wide dissection of globally emergent multi-drug resistant serotype 19A Streptococcus pneumoniae. BMC genomics, 2009. 10(1): p. 642.

24. Gherardi, G., et al., Serotype and clonal evolution of penicillin-nonsusceptible invasive Streptococcus pneumoniae in the 7-valent pneumococcal conjugate vaccine era in Italy. Antimicrobial agents and chemotherapy, 2012. 56(9): p. 4965–4968.

25. Dagan, R. and K.P. Klugman, Impact of conjugate pneumococcal vaccines on antibiotic resistance. The Lancet infectious diseases, 2008. 8(12): p. 785–795.

26. Farrell, D.J., K.P. Klugman, and M. Pichichero, Increased antimicrobial resistance among nonvaccine serotypes of Streptococcus pneumoniae in the pediatric population after the introduction of 7-valent pneumococcal vaccine in the United States. The Pediatric infectious disease journal, 2007. 26(2): p. 123–128.

27. Greene, S.K., et al., Trends in antibiotic use in Massachusetts children, 2000–2009. Pediatrics, 2012. 130(1): p. 15–22.

28. Pelton, S.I., et al., Emergence of 19A as virulent and multidrug resistant pneumococcus in Massachusetts following universal immunization of infants with pneumococcal conjugate vaccine. The Pediatric infectious disease journal, 2007. 26(6): p. 468–472.

29. Pichichero, M.E. and J.R. Casey, Emergence of a multiresistant serotype 19A pneumococcal strain not included in the 7-valent conjugate vaccine as an otopathogen in children. Jama, 2007. 298(15): p. 1772–1778.

30. Dagan, R., et al., Introduction and proliferation of multidrug-resistant Streptococcus pneumoniae serotype 19A clones that cause acute otitis media in an unvaccinated population. Journal of Infectious Diseases, 2009. 199(6): p. 776–785.

31. Gladstone, R.A., et al., Five winters of pneumococcal serotype replacement in UK carriage following PCV introduction. Vaccine, 2015. 33(17).

32. Diekmann, O. and J.A.P. Heesterbeek, *Mathematical epidemiology of infectious diseases*: model building, analysis and interpretation. Vol. 5. 2000: John Wiley & Sons.

33. Gupta, S., J. Swinton, and R.M. Anderson, Theoretical studies of the effects of heterogeneity in the parasite population on the transmission dynamics of malaria. Proceedings of the Royal Society of London B: Biological Sciences, 1994. 256(1347): p. 231–238.

34. Opatowski, L., et al., Contribution of mathematical modeling to the fight against bacterial antibiotic resistance. Current opinion in infectious diseases, 2011. 24(3): p. 279–287.

35. Caudill, L. and J.R. Wares, The role of mathematical modeling in designing and evaluating antimicrobial stewardship programs. Current Treatment Options in Infectious Diseases, 2016. 8(2): p. 124–138.

36. Obolski, U., G.Y. Stein, and L. Hadany, Antibiotic Restriction Might Facilitate the Emergence of Multi-drug Resistance. PLoS Comput Biol, 2015. 11(6): p. e1004340.

37. Obolski, U. and L. Hadany, Implications of stress-induced genetic variation for minimizing multidrug resistance in bacteria. BMC medicine, 2012. 10(1): p. 89.

38. Mitchell, P.K., M. Lipsitch, and W.P. Hanage, Carriage burden, multiple colonization and antibiotic pressure promote emergence of resistant vaccine escape pneumococci. Phil. Trans. R. Soc. B, 2015. 370(1670): p. 20140342.

39. Temime, L., D. Guillemot, and P. Boelle, Short-and long-term effects of pneumococcal conjugate vaccination of children on penicillin resistance. Antimicrobial agents and chemotherapy, 2004. 48(6): p. 2206–2213.

40. Lehtinen, S., et al., Evolution of antibiotic resistance is linked to any genetic mechanism affecting bacterial duration of carriage. Proceedings of the National Academy of Sciences, 2017. 114(5): p. 1075–1080.

41. Fish, D.N., S.C. Piscitelli, and L.H. Danziger, Development of resistance during antimicrobial therapy: a review of antibiotic classes and patient characteristics in 173 studies. Pharmacotherapy: The Journal of Human Pharmacology and Drug Therapy, 1995. 15(3): p. 279–291.

42. Gillespie, D.T., A general method for numerically simulating the stochastic time evolution of coupled chemical reactions. Journal of computational physics, 1976. 22(4): p. 403–434.

43. Jolley, K.A. and M.C. Maiden, BIGSdb: Scalable analysis of bacterial genome variation at the population level. BMC bioinformatics, 2010. 11(1): p. 1.

44. Corander, J. and P. Marttinen, Bayesian identification of admixture events using multilocus molecular markers. Molecular ecology, 2006. 15(10): p. 2833–2843.

45. Corander, J., et al., Enhanced Bayesian modelling in BAPS software for learning genetic structures of populations. BMC bioinformatics, 2008. 9(1): p. 1.

